# Molecular engineering of a cryptic epitope in Spike RBD improves manufacturability and neutralizing breadth against SARS-CoV-2 variants

**DOI:** 10.1101/2022.09.14.507842

**Authors:** Sergio A. Rodriguez-Aponte, Neil C. Dalvie, Ting Y. Wong, Ryan S. Johnston, Christopher A. Naranjo, Sakshi Bajoria, Ozan S. Kumru, Kawaljit Kaur, Brynnan P. Russ, Katherine S. Lee, Holly A. Cyphert, Mariette Barbier, Harish D. Rao, Meghraj P. Rajurkar, Rakesh R. Lothe, Umesh S. Shaligram, Saurabh Batwal, Rahul Chandrasekaran, Gaurav Nagar, Harry Kleanthous, Sumi Biswas, Justin R. Bevere, Sangeeta B. Joshi, David B. Volkin, F. Heath Damron, J. Christopher Love

**Author notes:** Contributed equally (Sergio A. Rodriguez-Aponte, Neil C. Dalvie, Ting Y. Wong). Present address: Merck Research Laboratories, West Point, PA 19486, USA.

## Abstract

There is a continued need for sarbecovirus vaccines that can be manufactured and distributed in low- and middle-income countries (LMICs). Subunit protein vaccines are manufactured at large scales at low costs, have less stringent temperature requirements for distribution in LMICs, and several candidates have shown protection against SARS-CoV-2. We previously reported an engineered variant of the SARS-CoV-2 Spike protein receptor binding domain antigen (RBD-L452K-F490W; RBD-J) with enhanced manufacturability and immunogenicity compared to the ancestral RBD. Here, we report a second-generation engineered RBD antigen (RBD-J6) with two additional mutations to a hydrophobic cryptic epitope in the RBD core, S383D and L518D, that further improved expression titers and biophysical stability. RBD-J6 retained binding affinity to human convalescent sera and to all tested neutralizing antibodies except antibodies that target the class IV epitope on the RBD core. K18-hACE2 transgenic mice immunized with three doses of a Beta variant of RBD-J6 displayed on a virus-like particle (VLP) generated neutralizing antibodies (nAb) to nine SARS-CoV-2 variants of concern at similar levels as two doses of Comirnaty. The vaccinated mice were also protected from challenge with Alpha or Beta SARS-CoV-2. This engineered antigen could be useful for modular RBD-based subunit vaccines to enhance manufacturability and global access, or for further development of variant-specific or broadly acting booster vaccines.

## Introduction

The global distribution of COVID-19 vaccines continues to lag in low- and middle-income countries (LMICs), which have struggled to acquire and distribute doses due to high prices and logistical distribution requirements such as cold chains^1^. Vaccines that effectively prevent severe COVID-19 symptoms and death are broadly available in high-income countries^2^, but SARS-CoV-2 variants of concern (VOCs) continue to emerge with increased transmissibility and the ability to escape known neutralizing antibodies^3–6^. Effective and accessible vaccines or boosters are still important, especially ones that can be manufactured at low costs, that can be stored at non-freezing or ambient temperatures, and could offer protection against VOCs.

Protein subunit-based vaccines typically can tolerate higher temperatures than mRNA vaccines for storage and transport and have been proven effective for prophylactic prevention of SARS-CoV-2^7,8^. While the SARS-CoV-2 spike (S) protein antigen elicits strong neutralizing responses when administered with adjuvants, the full trimeric form of the S protein remains difficult to manufacture^9,10^. Smaller subunit antigens such as the S protein receptor binding domain (RBD) also elicit protective immunity against SARS-CoV-2 in animal models and humans^11–15^. The RBD is also currently produced in microbial hosts with existing manufacturing infrastructure in LMICs^16,17^.

Improving the productivity of microbial fermentation can further lower the cost of manufacturing subunit vaccines^18^. Enhancements to stabilize subunit proteins can improve manufacturing yields^19^ and reduce costs associated with storage and distribution^20^. We previously used molecular engineering to improve the production of RBD in yeast, as well as its biophysical stability^11^. Here, we performed further molecular engineering of the SARS-CoV-2 RBD. We modified a hydrophobic region that is buried in the closed-form S protein, but is exposed on the RBD antigen itself. These changes led to a three-fold improvement in secreted titer and increased biophysical stability. The alterations described here did not significantly alter the antigenicity or immunogenicity of the RBD when displayed on a virus-like particle (VLP). VLP-RBD conjugates elicited immunity in mice against and protection against SARS-CoV-2 Alpha and Beta variants.

## Results

### Mutation of the RBD core hydrophobic patch to improve expression

We previously identified an exposed hydrophobic patch in the receptor binding motif (RBM) of the RBD that may cause both intracellular and extracellular aggregation of the RBD^11^. Two mutations to the RBM conferred a significant improvement to the production of secreted protein from yeast and the biophysical stability of the protein. These point mutations (L452K and F490W) were selected from the highly conserved sarbecovirus genetic background to retain binding to the human ACE2 receptor and minimize effects on antigenicity.

Here, we sought to apply similar rational engineering to a second predicted hydrophobic patch on the RBD core^21^, near the C-terminus. This hydrophobic patch is not predicted to be exposed in the native S protein with the RBD in the “down” confirmation (Fig. 1a). This patch is, however, exposed when the RBD is expressed independently as a soluble monomer. We hypothesized that, similar to the hydrophobic patch found in the RBM, the RBD core hydrophobic patch may reduce the solubility, stability, and efficient secretion of the RBD^22^. We sought here to eliminate or reduce this hydrophobic patch by rational substitutions of amino acids similar to our previous engineering efforts^11,16^.

**Fig. 1.**
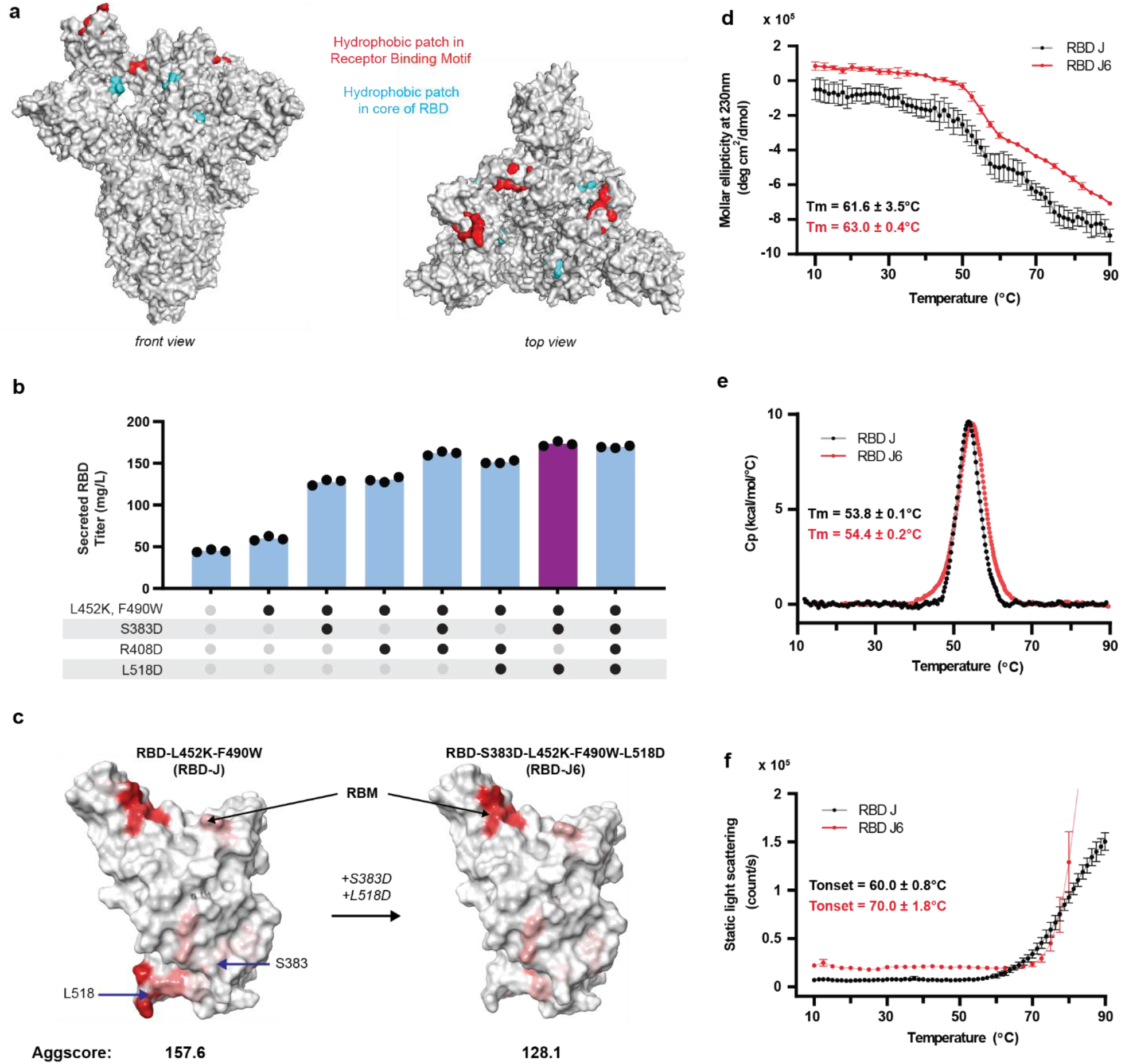
Molecular engineering of RBD core hydrophobic hotspot improves manufacturability. (**a**) Structual rendering of SARS-CoV-2 Spike trimer (PDB ID:7DK3). Amino acids included in RBM hydrophobic patch (red). Amino acids included in RBD core hydrophobic patch (blue). (**b**) Titer of mutated RBD secretion in 3mL plate cultures, measured by reverse-phase liquid chromatography. (**c**) Structural rendering of RBD-J and RBD-J6 with predicted hydrophobic patches (red). (**d**) Molar ellipticity at 230 nm as a function of increasing temperature of purified RBD-J and RBD-J6 as measured by Far-UV circular dichroism. Errors bars represent standard deviation of three independent measurements (**e**) Normalized DSC thermograms of purified RBD-J and RBD-J6. (**f**) Static light scattering as a function of increasing temperature of purified RBD-J and RBD-J6. The data in **e-f** are the mean of two independent experiments and the error bars in f represent the standard deviation.

We selected 21 mutations from a previously published deep mutational scanning analysis that appeared to boost expression in yeast while retaining binding to ACE2^23^. We tested each mutation individually with a 201 amino-acid RBD (amino acids 332-532 in the Spike protein). Each mutant RBD also included the mutation L452K, which, in our previous work, improved the expression and stability of the RBD^11^. We transformed each RBD variant into yeast and assessed the secretion of the RBD (Supplementary Fig. S1). As expected, the RBD variant with only the previously described L452K mutation was secreted with almost 60% higher titer than the Wuhan-Hu-1 RBD. Among RBD variants with additional mutations to RBD-L452K, we observed up to a ~4x improvement in specific productivity over the Wuhan-Hu-1 RBD. Interestingly, we found that three of the mutants that most improved secretion included mutations to an aspartic acid (S383D, R408D, and L518D) at locations in or near the RBD core hydrophobic patch (Supplementary Fig. S1). This result suggested that mutation of a heavily hydrophobic region may improve the secretion of this protein. Indeed, solvent-accessible hydrophobic regions can be destabilizing^24,25^, and partially unfolded complexes can promote aggregation^26^. Notably, aspartic acid substitutions have been shown to improve the solubility of antibody complementarity-determining regions (CDRs) ^27,28^.

We next evaluated combinations of the three aspartic acid substitutions in the RBD core hydrophobic patch. These designs also included both of the mutations to the hydrophobic patch in the RBM from our previous work (L452K, F490W, dubbed RBD-J)^11,29^. We observed the greatest improvement in the secreted titer (60mg/L to 173mg/L) upon addition of the S383D and L518D mutations (Fig. 1b). We computationally evaluated this variant of RBD-J and observed a reduction of the predicted surface hydrophobicity, AggScore^30^, from 157.6 to 128.1 (Fig. 1c). We concluded that this hybrid variant of RBD-J, S383D-L452K-F490W-L518D, denoted as RBD-J6, merited further characterization to assess its antigenicity and immunogenicity.

### Biophysical characterization of engineered SARS-CoV-2 RBDs

We compared the biophysical properties of RBD-J and RBD-J6. First, we performed far-UV circular dichroism (CD) spectroscopy as a function of temperature and observed that the thermal melting temperature (Tm) value of RBD-J6 (63.0 ± 0.4°C) was ~1.5°C higher than that of RBD-J (61.6 ± 3.5°C). This result suggested that the overall secondary structure of RBD-J6 was modestly more stable when compared to RBD-J (Fig. 1d). Next, we performed differential scanning calorimetry and observed that the Tm value of RBD-J6 (54.4 ± 0.1°C) was also slightly higher than the RBD-J (53.8 ± 0.1°C), a result consistent with the CD analysis, suggesting a small improvement to overall conformational stability of RBD-J6 (Fig. 1e). Lastly, we performed static light scattering as a function of temperature to assess the tendency of these two protein antigens to form aggregates. The thermal onset temperature (Tonset) value of RBD-J6 (70.0 ± 1.8°C) was ~10°C higher than that of RBD-J (60.0 ± 0.8°C) (Fig. 1f), suggesting the colloidal stability of the RBD-J6 was enhanced. Taken together, these results indicate that mutation of solvent accessible hydrophobic regions can improve overall conformational stability of the antigen, and consequently, colloidal stability as well.

### Antigenic characterization of engineered SARS-CoV-2 RBDs

Modification of the RBD to improve its physical qualities and stability as a vaccine candidate could potentially alter the antigenicity of the RBD. Unlike our previous engineering of the RBD, the two mutations introduced in RBD-J6 (S383D, L518D) are not native to the sarbecoviruses, and could affect the antigenicity of the RBD^31^. To assess the antigenicity of RBD-J6, we evaluated its binding to ACE2 and to several neutralizing antibodies targeting different epitopes of RBD^32,33^. We observed that RBD-J6 and RBD-J exhibited similar binding to ACE2 (Fig. 2a). RBD-J6 and RBD-J also exhibited similar binding to CV30^34^ and CB6^35^, two class I neutralizing antibodies that target the RBM (Fig. 2b, c), and to S309, a class III neutralizing antibody that targets the proteoglycan site of the RBD^36^ (Fig. 2d). We did observe, however, that RBD-J6 bound less strongly than RBD-J to CR3022 and EY6A, two class IV neutralizing antibodies that bind the RBD core near the C-terminus^33,35^ (Fig. 2e, f), suggesting that the modified residues altered the RBD core epitope created when expressing the RBD domain alone.

**Fig. 2.**
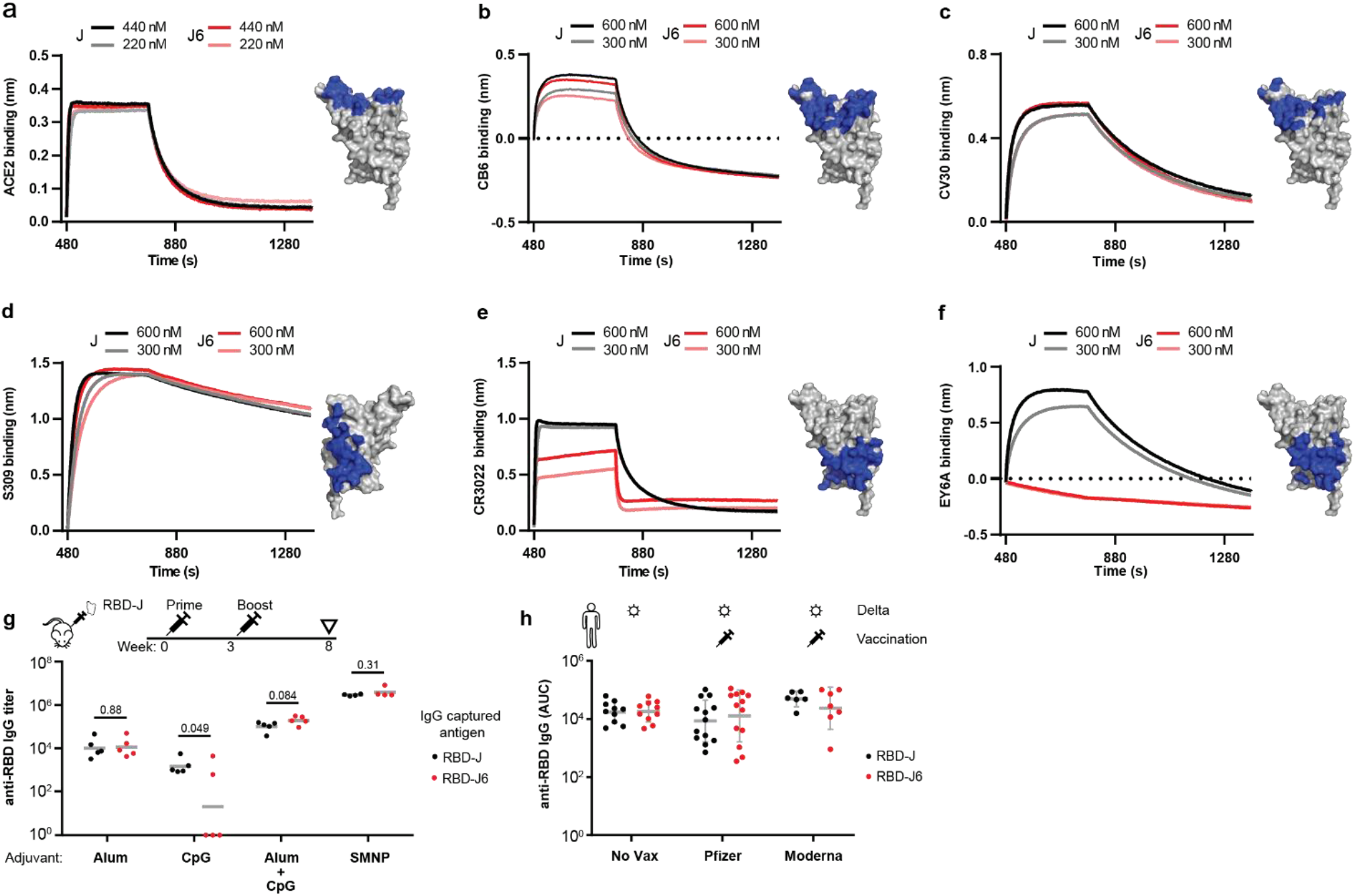
Antigenic comparison of RBD-J and RBD-J6. Binding of purified RBD to (**a**) human ACE2-Fc fusion protein, (**b**) CB6, class I neutralizing antibody, (**c**) CV30, class I neutralizing antibody, (**d**) S309, class III neutralizing antibody, (**e**) CR3022, class IV neutralizing antibody, and (**f**) EY6A, class IV neutralizing antibody by biolayer interferometry. Blue regions on the RBD structure indicate target binding epitopes. (**g**) Immunization timeline, titer of RBD-specific IgG of sera from RBD-J immunized mice with different adjuvants. Significance was determined by t-test. P-values are indicated in the plot. Gray bar represents mean values. (**h**) Area under the curve (AUC) for anti-RBD IgG binding titers of Delta variant convalescent sera from unvaccinated, Comirnaty (Pfizer) vaccinated, and Spikevax (Moderna) vaccinated cohorts. Gray bars represent mean and standard deviation of sample set.

We then evaluated how polyclonal antibodies raised in mice immunized with RBD-J bound to RBD-J6. In our previous study, we formulated RBD-J with different adjuvant combinations including alum alone, CpG1826 alone, alum + CpG1826, or a saponin-based adjuvant SMNP^11,37^. We found that antibodies in sera from mice immunized with alum alone, alum + CpG1826, or SMNP exhibited similar binding to both RBD-J and RBD-J6 (Fig. 2g).

These results suggest that antibodies raised against the adjuvanted RBD-J may not target the RBD core epitope when delivered with these adjuvants. Indeed, most potent neutralizing antibodies raised against the RBD target the RBM^38,39^. Interestingly, we observed that antibodies raised against RBD-J formulated with CpG1826 alone exhibited less binding to RBD-J6. Indeed, we have hypothesized previously that CpG adjuvants may alter the structure and antigenicity of soluble RBD^12^, and have recently demonstrated that CpG and aluminum salt adjuvants destabilize formulated RBD-J during storage^40^.

Finally, we evaluated the binding to both engineered RBDs of antibodies from convalescent sera obtained from patients infected with SARS-CoV-2 Delta (B.1.617.2), including ones whom had been vaccinated with approved mRNA COVID-19 vaccines before infection. The binding of these antibodies to either RBD-J or RBD-J6 was not significantly different, regardless of vaccination status (Fig. 2h, p = 0.7567, Paired t-test). These data, together, suggest that the S383D and L518D RBD mutations do not impact binding to a breadth of human antibodies raised in response to infection or vaccination where native S protein is present.

### Expression and antigenicity of RBD-J6 with mutations from circulating VOCs

Emerging variants of concern (VOCs) of SARS-CoV-2 have spurred the development of updated vaccine candidates that incorporate new viral mutations. In this study, we focused on two SARS-CoV-2 variants Alpha (B.1.1.7) and Beta (B.1.351). Several natural RBD mutations, including N501Y (Alpha and Beta), K417N (Beta), and E484K (Beta), reportedly increase virus transmissibility^41,42^ and evasion of neutralizing antibodies^43,44^. We sought to determine if the benefits in biophysical stability and manufacturability that we observed by mutation of the RBD core hydrophobic patch in RBD-J6 would also benefit RBD antigens that include mutations from the Alpha and Beta variants.

We added the three mutations found in the RBD of the Beta variant to RBD-J6 (hereafter, RBD-J6 β). We first evaluated the secreted expression of RBD-J6 β in yeast and observed 30% lower titers compared to RBD-J6. This reduced titer of the β version of RBD-J6 agrees with our previous report on the expression of the Wuhan RBD and its B.1.351 variant in yeast^16^. This reduced expression reduction also coincides with an increase of the AggScore from 128.1 to 216.5 when incorporating Beta variant mutations to RBD-J6 consistent with our hypotheses on effects of hydrophobicity on RBD stability.

We evaluated binding of RBD-J6 β to ACE2 and observed similar binding as RBD-J6 (Supplementary Fig. S2). Previous studies report that these mutations found in the RBD from SARS-CoV-2 Alpha and Beta increase affinity to ACE2^45^, but we did not observe this difference in combination with the RBD-J6 mutations. Both RBD-J6 and RBD-J6 β had similar binding to S309, CR3022 and EY6A antibodies as well (Supplementary Fig. S2). We observed that RBD-J6 β bound less strongly than RBD-J6 to the neutralizing antibodies CV30 and CB6, which target the RBM (Supplementary Fig. S2). This result suggests that these amino acids, K417, E484 and N501, are targeted by ACE2-blocking antibodies raised against the original Wuhan variant, and that the Beta variant mutations may reduce recognition of certain epitopes. With the exception of the class IV antibodies, RBD-J β exhibited a comparable binding profile to RBD-J6 β. Lastly, RBD-J β and RBD-J6 β were bound comparably by the same human convalescent sera from Delta SARS-CoV-2 breakthrough infections (Supplementary Fig. S2, p = 0.3214, Paired t-test).

### Vaccination of mice with RBD-J6 β and challenge with SARS-CoV-2 Alpha and Beta

We next sought to compare the immunogenicity and protectiveness of the RBD-J β and RBD-J6 β. We previously reported a vaccine design in which an RBD antigen was displayed on the surface of Hepatitis B surface antigen virus-like particles (HBsAg VLPs)^12^. In this design, attachment of the RBD to the HBsAg VLP was mediated by spontaneous conjugation of SpyTag and SpyCatcher peptides^46^. In addition to the demonstrated immunogenicity of this vaccine candidate in macaques rhesus^12^, the modular design of this system could allow integration of new RBD antigens such as RBD-J6 to improve stability and manufacturability, or RBD-J6 β to address new VOCs such as Beta. We chose, therefore, to evaluate the immunogenicity of RBD-J β and RBD-J6 β conjugated with HBsAg VLPs. RBD-J β and RBD-J6 β were expressed in yeast, and each purified RBD antigen was conjugated onto HBsAg VLPs (Supplementary Fig. S3). We evaluated the antigenicity of each vaccine and observed similar binding and avidity to ACE2 as well as S309 antibody for both VLP – RBD-J β conjugate and VLP – RBD-J6 β conjugate (Supplementary Fig. S3).

To evaluate the impact of the RBD-J6 mutations on the immunogenicity of RBD-based vaccines, we immunized K18-hACE2 transgenic mice^47–49^ intramuscularly with: 1) 10µg of VLP – RBD conjugate adjuvanted with 100µg of alum, or 2) 3µg of Comirnaty mRNA at weeks 0 and 3. At weeks 2, 5, and 7, we characterized the serological response against RBDs from several SARS-CoV-2 VOCs (Fig. 3a). After one dose, we observed higher RBD IgG titers across multiple VOCs in sera from mice immunized with the VLP – RBD-J6 β conjugate compared to sera from mice immunized with the VLP – RBD-J β conjugate (Fig. 3b). After three doses, the VLP – RBD-J6 β conjugate also elicited a higher RBD specific IgG response across multiple VOCs compared to the VLP – RBD-J β conjugate. These observed titers were similar to antibody titers elicited by two doses of Comirnaty, Pfizer-BioNTech’s mRNA vaccine (Fig. 3c).

**Fig. 3.**
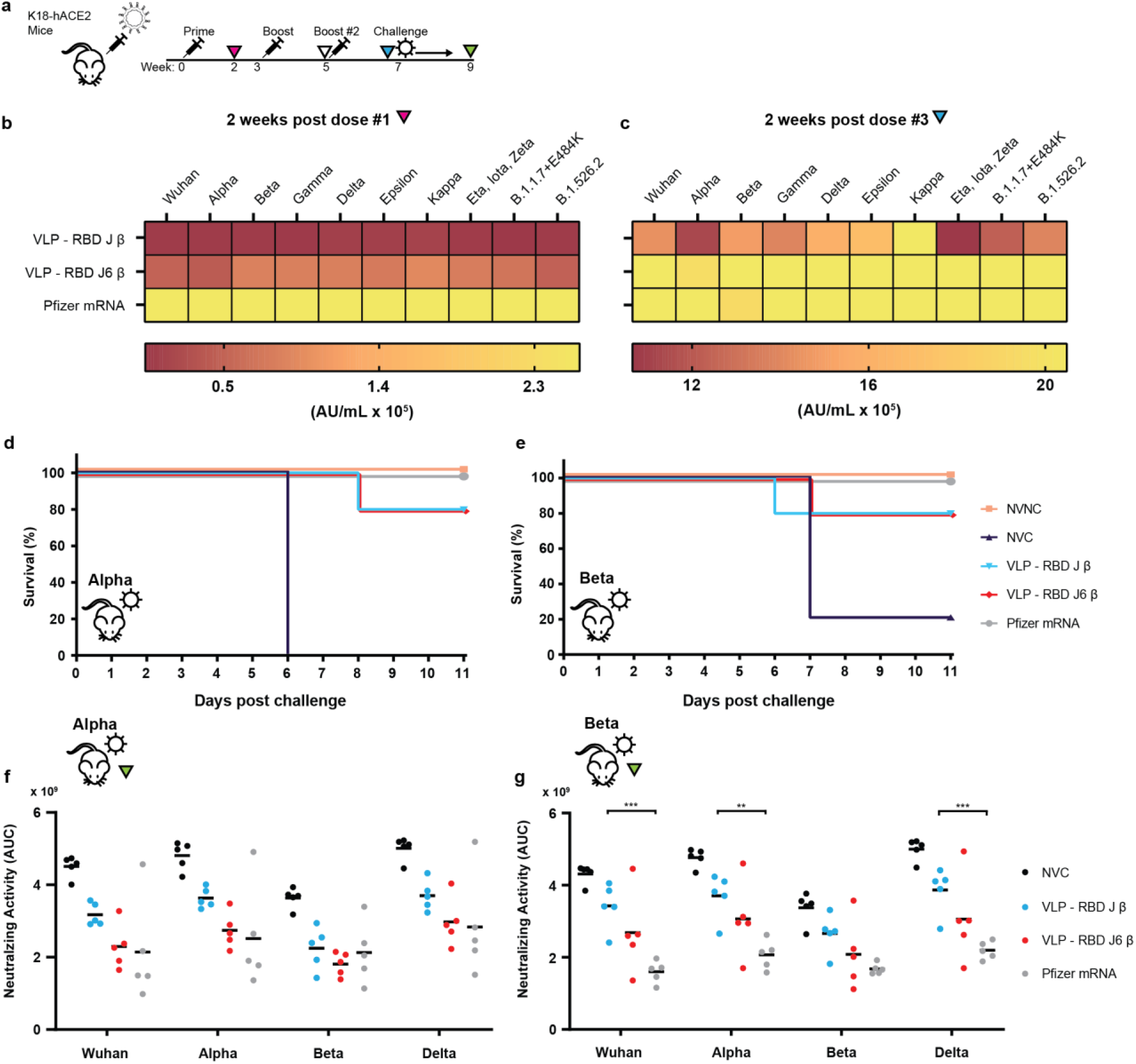
Immunogenicity and antigenicity of RBD-J6 β in K18-hACE2 mice challenged with SARS-CoV-2 Alpha or Beta variants. Vaccination and serology timepoint schedule for mice immunized with RBD conjugates (**a**). Heatmap of the mean IgG titer against SARS-CoV-2 VOC two weeks after prime dose (**b**) and four weeks after 2^nd^ dose (**c**). Kaplan-Meier survival curves for mice challenge with SARS-CoV-2 Alpha (**d**) and Beta (**e**). NVNC – non-vaccinated, non-challenged mice. NVC – non-vaccinated, challenged mice. MSD ACE2 neutralizing activity of VOC RBDs of post Alpha (**f**) and Beta (**g**) challenge sera against ancestral SARS-CoV-2 and variants. Points represent area under the curve of a serum dilution curve. Lower AUC indicates higher serum neutralizing activity. Black bars represent mean values. Significance was determined by two-way ANOVA using Tukey’s multiple comparison method (\**p*<0.05, \*\**p*<0.01, \*\*\**p*<0.001).

Next, at week 7, we challenged K18-hACE2 mice that received RBD-J β and RBD-J6 β vaccines with SARS-CoV-2 Alpha or Beta. We monitored weight loss and temperature for 11 days following challenge (Supplementary Fig. S4, S5). Four out of five mice immunized with VLP – RBD-J β or VLP – RBD-J6 β survived challenges from Alpha or Beta SARS-CoV-2, while non-vaccinated mice were morbid after Alpha (5/5) or Beta (4/5) challenges, respectively (Fig. 3d, e). All mice immunized with two doses of Comirnaty survived Alpha or Beta challenges. We evaluated the titers of viral RNA in the lung and brain of challenged mice. For both the Alpha and Beta challenge, we observed significantly reduced viral RNA in the lung and brain cells in mice immunized with either the VLP – RBD-J β or RBD-J6 β conjugates compared to the unvaccinated and challenged control group (Fig. 4). All detected viral RNA in mice immunized with VLP – RBD-J β or VLP – RBD-J6 β were statistically comparable to titers observed in mice that were immunized with Comirnaty, except in the lungs of mice challenged with SARS-CoV-2 Alpha. A histopathological analysis of pulmonary tissue showed reduced chronic and acute inflammation throughout the lung parenchyma after SARS-CoV-2 Alpha or Beta challenges compared to unvaccinated, challenged mice (Supplementary Fig. S6).

**Fig. 4.**
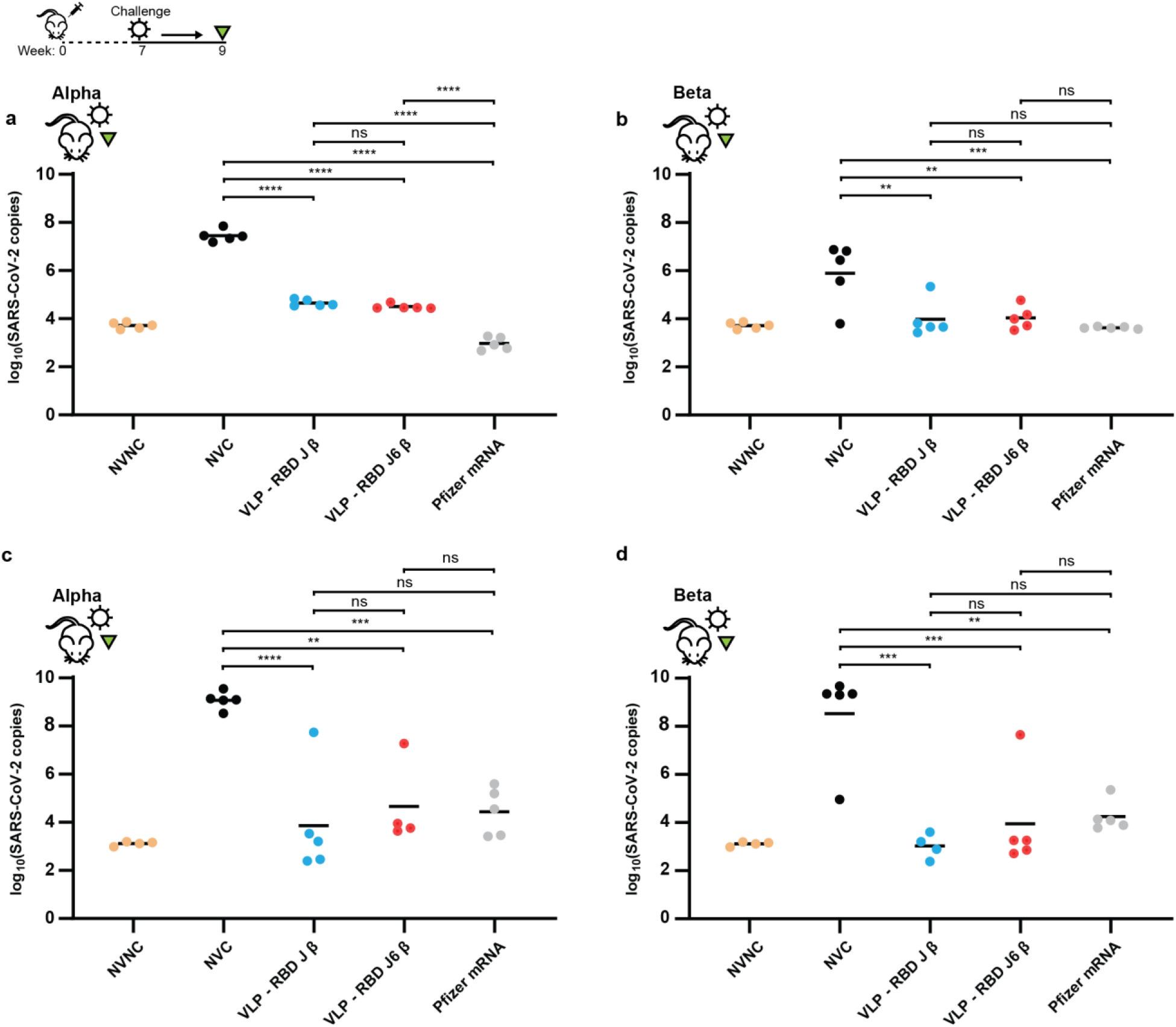
Viral RNA burden in lungs and brain of SARS-CoV-2 challenged mice. Nucleocapsid RNA copies in the lung right lobe of non-vaccinated, non-challenged (NVNC) mice, and mice challenged with SARS-CoV-2 Alpha (**a**), or Beta (**b**) variant. Nucleocapsid RNA copies in the brain of non-vaccinated, non-challenged (NVNC) mice, and mice challenged with SARS-CoV-2 Alpha (**c**), or Beta (**d**) variant. NVC – non-vaccinated, challenged. Black bars represent mean values. Significance was determined by One-way Ordinary ANOVA using Tukey’s multiple comparison method. \**p*<0.05, \*\**p*<0.01, \*\*\**p*<0.001, \*\*\*\**p*<0.001

Finally, we evaluated neutralizing activity of sera from mice challenged with Alpha and Beta variants against RBDs from SARS-CoV-2 VOC (in this assay, higher neutralizing activity is represented as lower area under the curve (AUC) of a dose-response curve between serum dilutions and electrochemiluminescence). No significant differences were observed in the sera neutralizing activity from mice immunized with either the VLP – RBD-J β or VLP – RBD-J6 β conjugates, and Comirnaty, after challenge with the Alpha variant (Fig. 3f). We also observed no significant differences in the neutralizing activity of sera from mice immunized with the VLP – RBD-J6 β conjugate and Comirnaty, after challenge with the Beta variant suggesting that the VLP – RBD-J6 β conjugate induces similar neutralizing potency across VOC RBDs compared to Comirnaty (Fig. 3g). Sera from mice immunized with the VLP – RBD-J β conjugate and challenged with Beta, however, exhibited significantly lower neutralization of Wuhan, Alpha and Delta variants compared to sera from mice immunized with Comirnaty (Fig. 3g).

## Discussion

We previously described engineering the SARS-CoV-2 RBD to improve manufacturability, stability, and immunogenicity. Specifically, we reduced the predicted hydrophobicity of the receptor binding motif (RBM) in the engineered variant RBD-J^11^. Here, we report a second generation engineered variant, RBD-J6 (RBD-S383D-L452K-F490W-L518D), which further improves manufacturability and stability due to two mutations that reduce the predicted hydrophobicity of a second hydrophobic patch on the core of the RBD subunit.

Notably, *in situ*, this second hydrophobic patch is primarily exposed when the full S protein RBD is in the “up” position (Fig. 1a), and residues like S383 form hydrogen bonds with surrounding S trimer domains^50,51^. This report agrees with other studies that have also introduced mutations near the RBD core, leading to increased production and stability^52,53^. These results and the general strategy used may inform approaches to engineer other vaccine antigens and therapeutic proteins based on protein subunits that create new surface exposed hydrophobic patches when truncated from their full-length protein. Monoclonal antibodies are often designed and selected for reduced surface hydrophobicity during development, which has been shown to be a critical factor associated with clinical success^54,55^.

We also evaluated the antigenicity of the RBD-J6 β variant and observed that binding to the target receptor ACE2 and several known neutralizing antibodies was not impacted by the addition of mutations in the RBD core hydrophobic patch. Binding was only disrupted, however, for antibodies that targeted the mutated epitope. Notably, these antibodies that target the class IV epitope do not block binding of the RBD to the ACE2 receptor. These results suggest that the overall secondary and tertiary structure of the protein was not measurably impacted by the additional mutations. When we tested the RBD-J6 antigen in a transgenic mouse model, we observed higher titers of antibodies that bind to several RBD variants compared to the RBD-J antigen (Fig. 3). Likewise, the RBD-J6 antigen elicited improved neutralizing activity of several SARS-CoV-2 VOC RBDs in comparison to the RBD-J antigen (Fig. 3). This result demonstrates that the immunogenicity and antigenicity of the RBD-J6 antigen are sufficient, despite modification of an epitope for previously identified neutralizing antibodies^51^. We hypothesize, furthermore, that the apparent breadth of antibody binding across RBD variants conferred by immunization with the RBD-J6 antigen may be due to focusing of the immune response on other, possibly broadly neutralizing epitopes^56^. The breadth that could be conferred by focusing on highly conserved epitopes is an active field of research with respect to SARS-CoV antigens and other pathogens^57–59^.

Over two rounds of rational engineering, we have now achieved a >10-fold increase in expression titer over the sequences for the ancestral RBD, and overall improvements to its biophysical stability^11^. Surprisingly, both rounds of rational engineering also improved the potency and breadth of the immune response to the RBD antigen compared to the native ancestral sequence. Together, these improvements suggest a correlation exists among protein structure, biophysical properties, and antigenic response. Indeed, stabilization of protein-based vaccines has been reported to enhance MHC presentation and TH1 responses^60^.

Mice challenged with Alpha and Beta variants of SARS-CoV-2 after immunization with protein subunit VLP-RBD conjugates had nearly complete protection, and the neutralizing activity elicited by three doses of the VLP – RBD-J6 β conjugate was comparable to two doses of Comirnaty. Recent literature has demonstrated that individuals immunized with currently approved mRNA vaccines show a marked decline in anti-RBD IgG titers after several months^61,62^, suggesting that vaccine boosters may be needed to maintain long-term immunity^63^. Several studies have reported using RBD as a booster for Spike protein primed animals^64,65^.

Future studies are merited to evaluate the use of this VLP-RBD candidate or other multimeric versions as a 1st or 2nd booster to an mRNA vaccination. We also hypothesize that ablation of the RBD core epitope targeted by class IV antibodies in this RBD candidate could focus an existing immune memory to more variable RBM epitopes with higher neutralizing rates^38,39^.

To date, the optimal frequency of vaccine boosters for SARS-CoV-2 and, subsequently, the overall yearly global demand for vaccine doses are unknown^63^. Despite the increased access of boosters in developed countries, likely tens of billions of vaccine doses will need to be manufactured and distributed at low cost to maintain global long-term immunity, suppress the formation and spread of new VOCs, and reach all populations with limited access to healthcare^66,67^. We have demonstrated that the improved RBD antigen reported here is compatible with several vaccines designed for access in low- and middle-income countries^16^.

We believe that the improved manufacturability, breadth, and stability of RBD-J6 can be translated to RBD-based clinical candidates like CorbeVax™^68^, Soberana, Cuba’s RBD conjugate vaccine^69^, and others^70–72^. The enhancements to the RBD presented here are an important step towards accessible, low-cost, and effective RBD-based COVID vaccines^73^.

## Materials and Methods

### Strains

Recombinant genes for RBD variant expression were codon optimized for *Komagataella phaffii* expression, and synthesized cloned into a custom vector on a BioXP (Codex). Linear DNA was purified, and constructs were transformed into a modified wild-type *K. phaffii* (NRRL Y-11430) as described previously^11,74^.

### Cultivations

Strains for titer evaluation of each RBD mutant were cultivated in 3mL in 24-deep well plates as described previously^11^. Strains producing material for further analytical characterization and mice immunization were cultivated in 200mL in 1L baffled flasks as described previously^11^. Cells were inoculated at 0.1 OD_600_, growth for 24 h in complex media containing 4% glycerol, pelleted, resuspended, and grown for 24 h in complex media containing 40 g/L sorbitol and 1% methanol.

### Protein Purification

Purification of harvested cell culture supernatant containing the recombinant protein was performed using the downstream processing (DSP) module of InSCyT as described previously^11,75^. The supernatant pH was adjusted inline to pH 5.0 with 100mM citric acid. The supernatant was loaded into a 5mL prepacked CMM HyperCel column (Sartorius AG, Gottingen, Germany). The column was re-equilibrated with 20mM citric acid, pH 5.0, washed with 20mM sodium phosphate pH 6.5, and elution was initiated with 20mM sodium phosphate, 300mM NaCl, pH 8.5. Eluate from column 1 above 20mAu was loaded into a 1mL prepacked HyperCel STAR AX column (Sartorius AG, Gottingen, Germany). Flow-through material above 20mAu was collected.

### Preparation of vaccine materials

Purified RBD-SpyTag was conjugated onto Hepatitis B surface antigen (HBsAg) SpyCatcher VLPs overnight at a 1.5:1 RBD to HBsAg molar ratio. Conjugated VLP-RBD was buffer exchanged and concentrated with 100K molecular weight cutoff centrifugal filters.

Materials were formulated with a 10mM Histidine, 20mM Sodium phosphate, 5mM Tris, 37.5mM NaCl, pH 7.4 buffer. VLP-RBD formulations were diluted to a final concentration of 100ug/mL. Alhydrogel (Invivogen) was added to a final concentration of 400ug/mL.

### Reverse phase chromatography

Reverse phase high performance liquid chromatography (HPLC) for supernatant titer measurement was performed on an Agilent 1260 HPLC system (Agilent Technologies, Santa Clara, CA) using a PLRP-S column (2.1 x 150 mm, 300Å, 3µm) (Agilent Technologies, Santa Clara, CA) as described previously^11^.

### Far-UV Circular Dichroism (CD) spectroscopy

Far-UV CD spectroscopy was performed using a Chirascan-plus CD spectrometer (Applied Photophysics Ltd., Leatherhead, UK) equipped with a 6-cuvette position Peltier temperature controller (Quantum Northwest, Liberty Lake, WA) and a high-performance solid-state detector as described previously^11^.

### Static Light Scattering (SLS)

SLS measurements as a function of temperature were made in triplicate using a dual emission PTI QM-40 Spectrofluorometer (Horiba Scientific Northampton, UK) equipped with a 4-position cell holder Peltier temperature control device, a high-power continuous 75 W short-arc Xe lamp (Ushio), and a Hamamatsu R1527 photomultiplier tube as described previously^11^.

### Differential Scanning Calorimetry (DSC)

DSC was performed in duplicate using an auto-VP capillary differential scanning calorimeter (MicroCal/GE Health Sciences, Pittsburgh, PA) equipped with Tantalum sample and reference cells pressurized at ~60 psi with nitrogen as described previously^11^.

### Biolayer interferometry (BLI)

Biolayer interferometry was performed using the Octet Red96 with Protein A (ProA) biosensors (Sartorius ForteBio, Fremont, CA), which were hydrated for 15 min in kinetics buffer prior to each run. Kinetics buffer comprising 1X PBS pH 7.2, 0.5% BSA, and 0.05% Tween 20 was used for all dilutions, baseline, and disassociation steps. ACE2-Fc was loaded at a concentration of 10 ug/mL. Antibodies CB6, CV30, S309, CR3022, EY6A were loaded at a concentration of 2ug/mL. Samples were loaded in a 96-well black microplate (Greiner Bio-One, Monroe, NC) at starting concentrations of 15 µg/mL for ACE2-Fc binding and 10 µg/mL for antibody binding. Association and dissociation were measured at 1000 rpm for 300 and 600 seconds, respectively. Binding affinity was calculated using the Octet Data Analysis software v10.0 (Pall ForteBio), using reference subtraction, baseline alignment, inter-step correction, Savitzky-Golay filtering, and a global 1:1 binding model.

### Animal welfare, Biosafety and Ethics statements

B6.Cg-Tg(K18-ACE2)2Prlmn/J mouse vaccine and SARS-CoV-2 challenge studies were executed under IACUC protocol number 2009036460. All mice were humanely euthanized based on the disease scoring system^49^, and no deaths occurred in the cage. All SARS-CoV-2 challenge studies were conducted in the West Virginia University Biosafety Laboratory Level 3 facility under the IBC protocol number 20-04-01. SARS-CoV-2 samples were either inactivated with 1% Triton per volume or Trizol before exiting high containment.

### Mice immunization

Four week old B6.Cg-Tg(K18-ACE2)2Prlmn/J mice were purchased from Jackson Laboratory (stock no: 034860). Mice were first primed with the VLP-RBD vaccine at 9 weeks old, and boosted 3 and 6 weeks after. Mice immunized with Comirnaty were primed at 9 weeks old and boosted 3 weeks later. Mice were immunized intramuscularly with 50uL of vaccine formulation.

### MSD serological assay for IgG titer measurement

Anti-RBD IgG was measured in mice serum on a MSD QuickPlex SQ120 following the SARS-CoV-2 Plate 11 Multi-Spot 96-well, 10 spot plate manufacturer’s protocol. Sera from unvaccinated mice collected at weeks 2 and 7 was diluted 1:1000. Sera from vaccinated mice collected two weeks post prime (week 2) was diluted 1:4000-1:512000, and vaccinated sera collected two weeks post 2nd booster (week 7) was diluted 1:32000-1:4096000.

### MSD COVID-19 ACE2 neutralization assay

SARS-CoV-2 challenged serum was analyzed using the SARS-CoV-2 Plate 11 Multi-Spot 96-well, 10 spot plate following the manufacturer protocol (catalog #: K15458U-2) on the MSD QuickPlex SQ120. The 10 spots contained the following RBD antigens, common designations, and lineages: 1) Epsilon - L452R (B.1.427; B.1.429; B.1.526.1) 2) Beta - K417N, E484K, N501Y (B.1.351; B.1.351.1) 3) Eta, Iota, Zeta - E484K (B.1.525; B.1.526; B.1.618; P.2; R.1) 4) Gamma - K417T, E484K, N501Y (P.1) 5) New York - S477N 6) Alpha - N501Y (B.1.1.7) 7) UK, Philippines - E484K, N501Y (B.1.1.7+E484K; P.3) 8) Kappa - L452R, E484Q (B.1.617; B.1.617.1; B.1.617.3) 9) Delta - L452R, T478K (AY.3; AY.4; AY.4.2; AY.5; AY.6; AY.7; AY.12; AY.14; B.1.617.2; B.1.617.2+Δ144) and 10) Wuhan. Four dilutions of serum, 1:5, 1:50, 1:500, and 1:5000 was analyzed on the MSD neutralization assay for each mouse to perform Area Under the Curve analysis on the electrochemiluminescence using GraphPad Prism.

### SARS-CoV-2 propagation and mouse challenge

Alpha (NR-54000) and Beta (NR-54008) SARS-CoV-2 variants were obtained from BEI Resources. Alpha and Beta VOC were propagated in Vero E6 cells (ATCC-CRL-1586) and resequenced before use in mouse challenge. K18-hACE2 mice were anesthetized using an intraperitoneal injection of ketamine (Patterson Veterinary 07-803-6637, 80 mg/kg) / xylazine (07-808-1947, 8.3 mg/kg) and were intranasally challenged with 50uL of 104 PFU/dose of Alpha or Beta variant, 25uL per nare. Mice were monitored until awake.

### Disease monitoring of SARS-CoV-2 challenged mice

Challenged K18-hACE2 mice were evaluated daily through both in-person health assessments in the BSL3 and SwifTAG Systems video monitoring for 11 days. Disease assessments of the mice were scored based on five criteria: 1) weight loss (scale 0-5), 2) appearance (scale 0-2), 3) activity (scale 0-3), 4) eye closure (scale 0-2), and 5) respiration (scale 0-2)^49^. All five criteria were scored based off a scaling system where 0 represents no symptoms and the highest number on the scale denotes the most severe disease phenotypes. Additive disease scores of the five criteria were assigned to each mouse after evaluation. Mice that scored an additive disease score of 5 or above among all 5 criteria, or weight loss of 20% or greater during the disease assessment required immediate euthanasia. Cumulative disease scoring was calculated by adding the disease scores of each mouse from each group. Morbid mice that were euthanized during the study, before day 11, retained their disease score for the remainder of the experiment^76^.

### Euthanasia and tissue collection

Challenged mice that were assigned a disease score of 5 or above or reached the end of the experiment were euthanized with an IP injection of Euthasol (390mg/kg) (Pentobarbital) followed by secondary measure of euthanasia with cardiac puncture. Blood from cardiac puncture was collected in BD Microtainer gold serum separator tubes (BD 365967), centrifuged at 15,000 x g for 5 minutes and serum was collected for downstream analysis. Lungs were separated into right and left lobes. Right lobe of the lung was homogenized in 1mL of PBS in gentleMACS C tubes (order number: 130-096-334) using the m_lung_02 program on the gentleMACS Dissociator. 300μL of lung homogenate was added to 1000μL of TRI Reagent (Zymo research) for downstream RNA purification and 300μL of lung homogenate was centrifuged at 15,000 x g for 5 minutes and the lung supernatant was collected for downstream analyses. Brain was excised from the skull and was homogenized in 1mL PBS in gentleMACS C tubes using the same setting as lung on the gentleMACS Dissociator. 500μL of TRI Reagent was added to 1000μL of brain homogenate for RNA purification.

### SARS-CoV-2 viral RNA analysis of lung and brain by q-RT-PCR

RNA purification of the lung and brain were performed using the Direct-zol RNA miniprep kit (Zymo Research R2053) following the manufacturer protocol. SARS-CoV-2 copy numbers were assessed through qPCR using the Applied Biosystems TaqMan RNA to CT One Step Kit (Ref: 4392938), as described previously^76^.

### Lung Histopathology

Left lobes of lungs were fixed in 10 mL of 10% neutral buffered formalin. Fixed lungs were paraffin embedded into 5 μm sections. Sections were stained with hematoxylin and eosin (H&E) and were analyzed by iHisto. Lungs were scored by a pathologist for chronic and acute inflammation in the lung parenchyma, blood vessels, and airways, as described previously^76^.

## Acknowledgements

This work was funded by the Bill and Melinda Gates Foundation (Investment IDs INV-002740). This study was also supported in part by the Koch Institute Support (core) Grant P30-CA14051 from the National Cancer Institute. N.C.D. is supported by a graduate fellowship from the Ludwig Center at MIT's Koch Institute. The content is solely the responsibility of the authors and does not necessarily represent the official views of the NCI, the NIH, Gates MRI, or the Bill and Melinda Gates Foundation.

## Author contributions

S.R.A., N.C.D., and J.C.L. conceived and planned experiments. N.C.D. and R.S.J. generated and characterized yeast strains. S.R.A. and C.A.N performed HPLC assays. S.R.A. designed and performed protein purifications. S.B., K.K. and O. K. performed biophysical characterization. S.R.A performed antigenic characterization by BLI. H.D.R, M.P.R, R.R.L., U.S.S, S.B., R.C., G.N., and M.R. produced, purified and validated the HBsAg VLP material. N.C.D. and S.R.A. formulated and analyzed samples for animal studies. T.Y.W., B.P.R., K.S.L., H.A.C., and M.B. designed and performed mice immunizations, challenge and monitoring. T.Y.W. performed RBD ELISA assays. S.R.A., N.C.D., T.Y.W. and J.C.L. wrote the manuscript. J.C.L., F.H.D.., D.B.V., S.B.J., J.R.B., S.B, and H.K. designed the experimental strategy and reviewed analyses of data. All authors reviewed the manuscript.

## Competing interests

S.R.A., N.C.D., and J.C.L. have filed a patent related to the RBD-L452K-F490W (RBD-J) sequence. J.C.L. has interests in Sunflower Therapeutics PBC, Honeycomb Biotechnologies, OneCyte Biotechnologies, QuantumCyte, Amgen, and Repligen. J.C.L.’s interests are reviewed and managed under MIT’s policies for potential conflicts of interest. H.D.R, M.P.R, R.R.L., U.S.S., S.B., R.C., and G.N. are employees of Serum Institute of India Pvt. Ltd.

## Data Availability

All data generated or analyzed during this study are included in this published article (and its supplementary information files).

---

**Supplementary Fig. S1.**
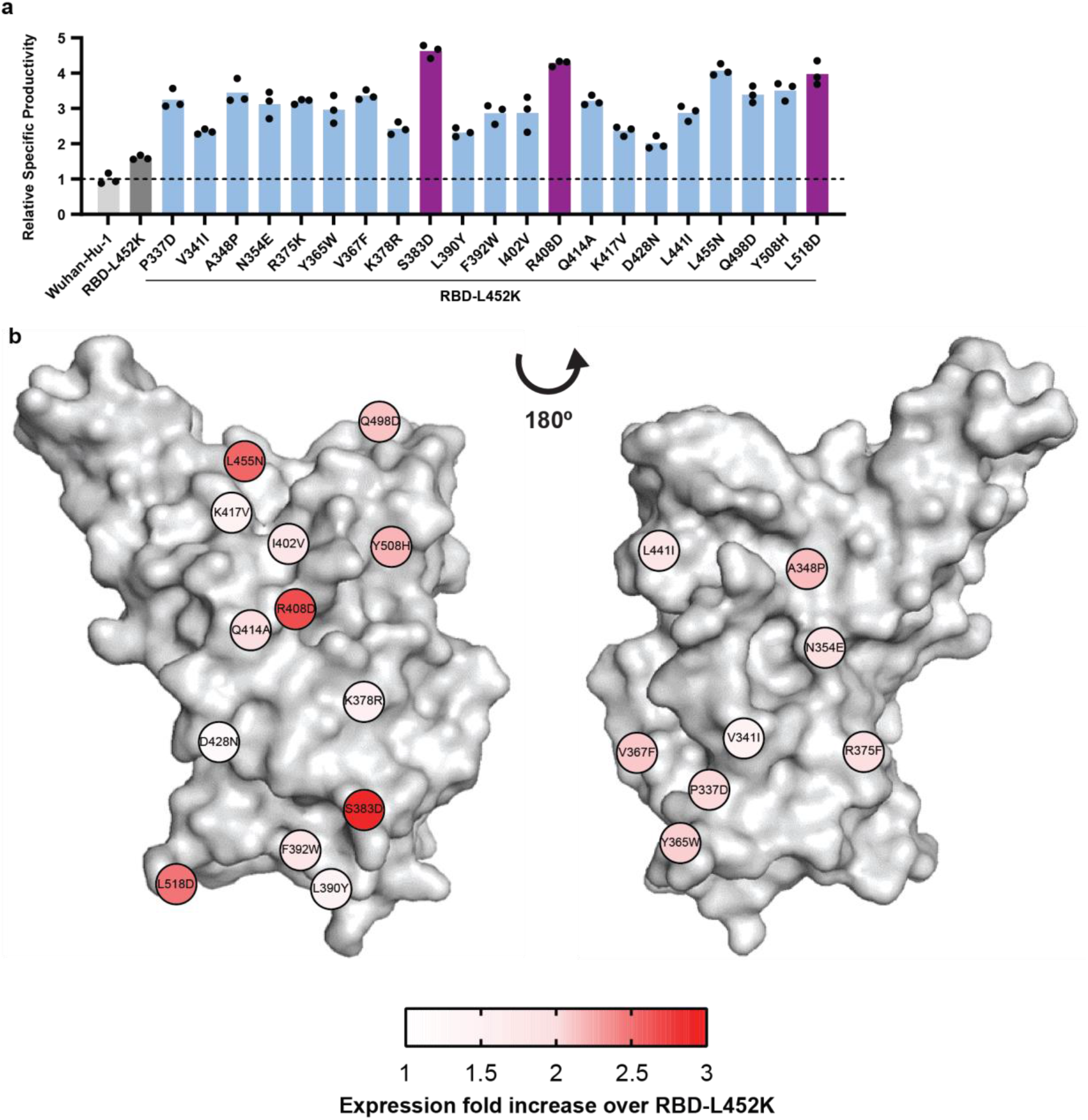
Expression titer of 21 mutations coupled with RBD-L452K. (**a**) Titer of mutated RBD secretion in 3mL plate cultures, measured by reverse-phase liquid chromatography. Bars represent mean values. (**b**) Localization of mutations coupled with RBD-L452K on RBD surface. Mutations are color-coded according to their expression fold increase over RBD-L452K.

**Supplementary Fig. S2.**
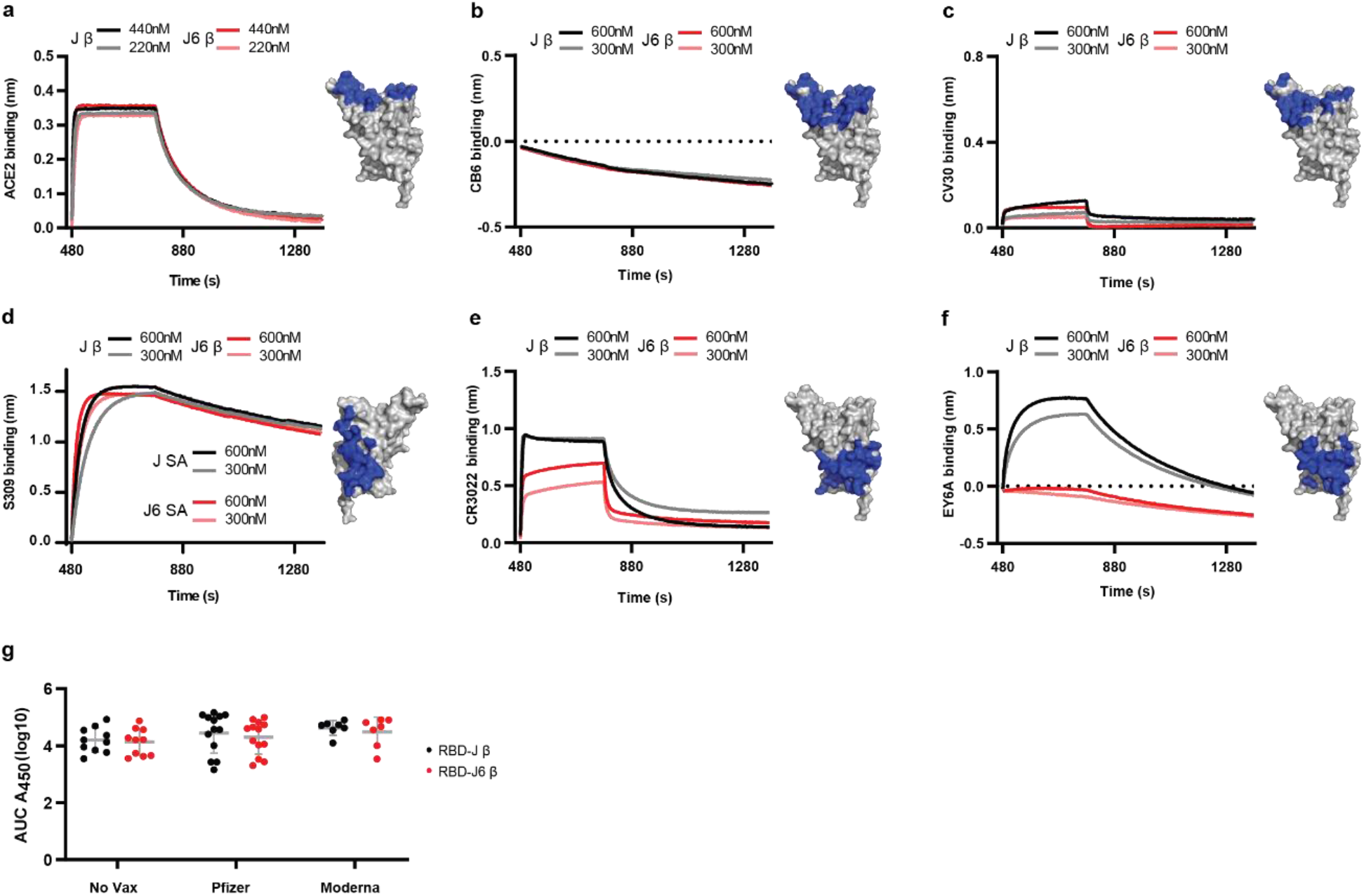
Antigenic characterization of RBD-J β and RBD-J6 β. Binding of purified RBD to (**a**) human ACE2-Fc fusion protein, (**b**) CB6, class I neutralizing antibody, (**c**) CV30, class I neutralizing antibody, (**d**) S309, class III neutralizing antibody, (**e**) CR3022, class IV neutralizing antibody, and (**f**) EY6A, class IV neutralizing antibody by biolayer interferometry. Blue regions on the RBD structure indicate target binding epitopes. (**g**) Area under the curve for antibody binding titers of Delta variant breakthrough cases’ convalescent sera from unvaccinated, Comirnaty (Pfizer) vaccinated, and Spikevax (Moderna) vaccinated cohorts. Gray bars represent mean and standard deviation of sample set.

**Supplementary Fig. S3.**
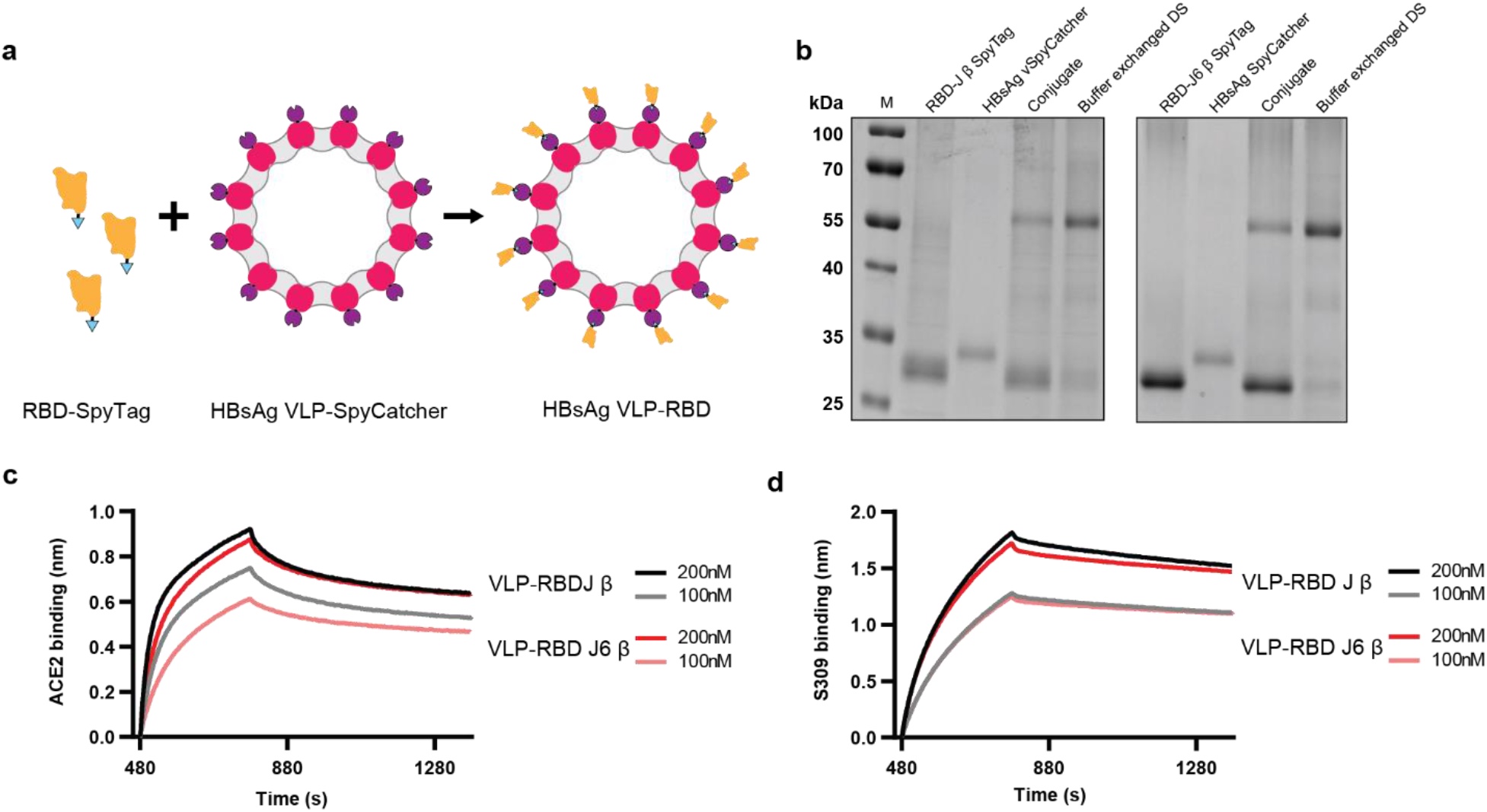
Design and characterization of VLP-RBD drug product. (**a**) Schematic of RBD-SpyTag conjugation onto HBsAg-SpyCatcher VLP. (b) Reduced SDS-PAGE analysis of conjugated RBD-J β VLP (left) and RBD-J6 β VLP (right). (**c-d**) Binding of VLP-RBD to (**c**) human ACE2-Fc fusion protein and (**d**) S309, class III neutralizing antibody, measured by biolayer interferometry.

**Supplementary Fig. S4.**
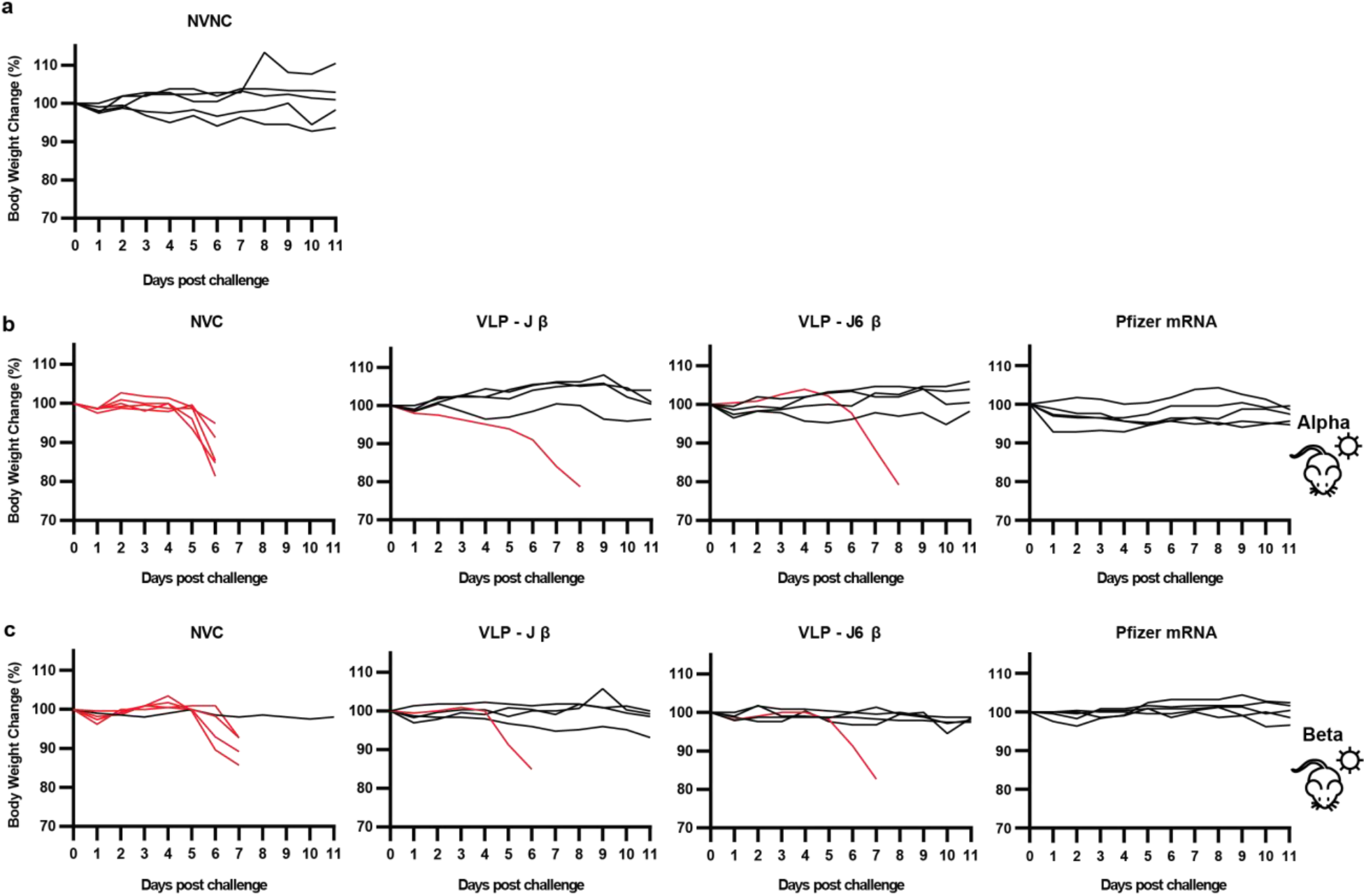
Body weight change during SARS-CoV-2 challenge. Body weight tracking of (**a**) non-vaccinated, non-challenged (NVNC) mice, (**b**) mice challenged with SARS-CoV-2 Alpha variant, and (**c**) mice challenged with SARS-CoV-2 Beta variant. Body weight of euthanized mice are labeled in red. NVC – non-vaccinated, challenged mice.

**Supplementary Fig. S5.**
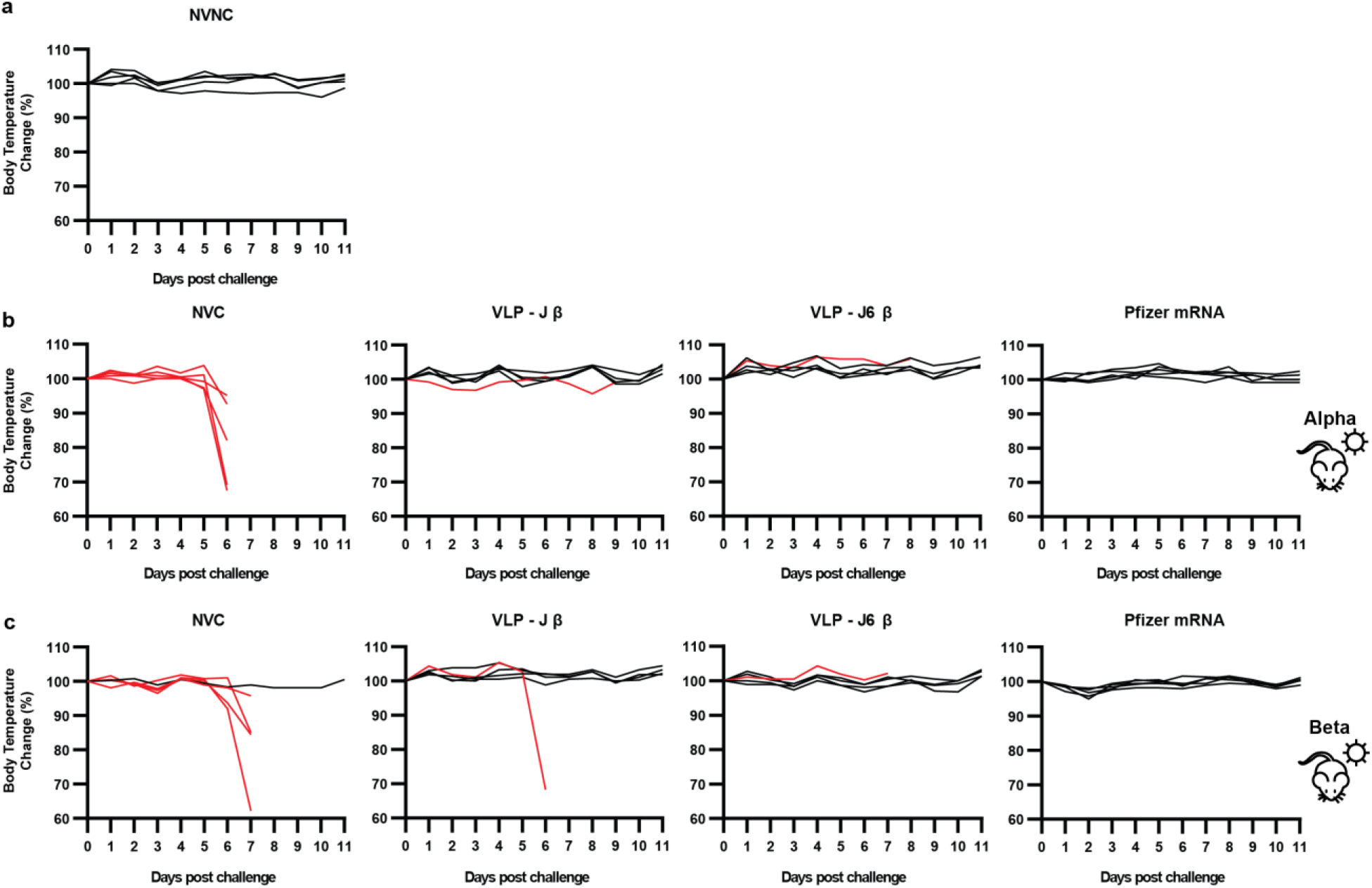
Body temperature change during SARS-CoV-2 challenge. Body temperature tracking of (**a**) non-vaccinated, non-challenged (NVNC) mice, (**b**) mice challenged with SARS-CoV-2 Alpha variant, and (**c**) mice challenged with SARS-CoV-2 Beta variant. Body temperature of euthanized mice are labeled in red. NVC – non-vaccinated, challenged mice.

**Supplementary Fig. S6.**
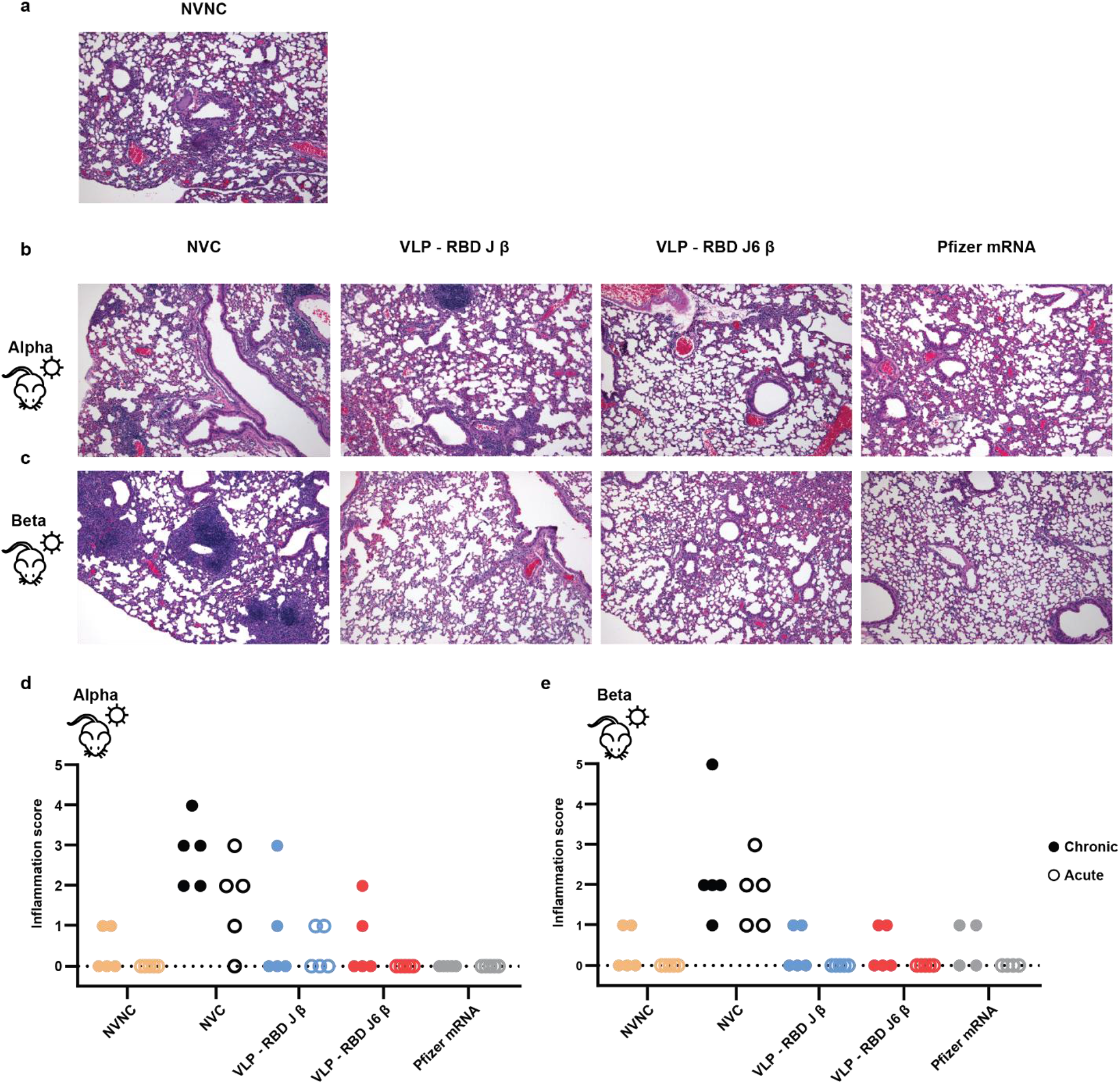
Histopathological analysis of lung tissue from SARS-CoV-2 challenged mice. Hematoxylin and eosin staining of lung tissue from non-vaccinated, non-challenged (NVNC) (**a**), SARS-CoV-2 Alpha (**b**) and Beta (**c**) challenged mice. Images presented at a 100x magnification. Chronic and acute inflammation scores of lungs from NVNC mice and SARS-CoV-2 Alpha (**d**) and Beta (**e**) variant challenged mice.

## Notes

### Competing Interest Statement

Sergio A. Rodriguez Aponte, Neil C. Dalvie., J. Christopher Love have filed a patent related to the RBD-L452K-F490W (RBD-J) sequence. J. Christopher Love has interests in Sunflower Therapeutics PBC, Honeycomb Biotechnologies, OneCyte Biotechnologies, QuantumCyte, Amgen, and Repligen. J. Christopher Love interests are reviewed and managed under MIT policies for potential conflicts of interest. Harish D. Rao, Meghraj P. Rajurkar, Rakesh R. Lothe, Umesh S. Shaligram, Saurabh Batwal, Rahul Chandrasekaran, and Gaurav Nagar are employees of Serum Institute of India Pvt. Ltd.

